# The Asymmetrical Architecture of Ultradian REM Sleep: Direction, NREM Currency, and a Finite Post-REM Predictive Horizon in Mice

**DOI:** 10.64898/2026.09.08.750050

**Authors:** Olivier Le Bon

**Author notes:** Correspondence: Olivier Le Bon.

## Abstract

Two accounts published in 1994 placed the controlling variable of the ultradian NREM-REM cycle on opposite sides of the REM episode: propensity accumulating during NREM sleep (Benington and Heller, 1994), or a constraint imposed by the preceding episode (Vivaldi et al., 1994). The Asymmetrical Hypothesis (Le Bon, 2021) proposed a third position, in which the preceding episode determines only a bounded post-REM refractory period rather than the whole interval. I tested these accounts in 1,955 valid inter-REM intervals reconstructed from continuous 24-hour recordings in 25 mice of three strains. Accumulated NREM did not predict the duration of the following REM episode (Spearman rho = 0.015, p = 0.50), whereas preceding REM duration predicted subsequent NREM accumulation (rho = 0.506) in every animal considered individually; a mixed-effects model fitted on the log-log scale returned an elasticity of 0.56 ± 0.03, steeper than previously reported log-log slopes near 0.33. A lower envelope near 1 min of accumulated NREM per minute of preceding REM survived leave-one-mouse-out validation and within-mouse permutation (6.1% of held-out intervals crossed it, against 10.2% under permutation), but was convex rather than proportional to the preceding episode. The influence of the preceding episode became undetectable beyond approximately 10 min of accumulated NREM, which outperformed elapsed chronological time as the relevant clock in held-out mice. Episode-termination hazard rose with elapsed episode duration (Weibull shape 1.48, versus 0.84 for inter-REM intervals), and inter-animal variability in episode number exceeded that in mean episode duration approximately fourfold. Part of the non-monotonic, rise-and-fall shape reported in recent propensity analyses arises from survivor selection rather than from any additional mechanism: intervals still at risk late are enriched in long preceding episodes, and a mixture of exponential hazards with no individual decline reproduces the same pooled pattern. Together, these findings indicate that the early post-REM constraint is real but neither strictly bounded nor proportional, expiring within about ten minutes rather than governing the whole interval; because episode duration is itself internally constrained, 24-hour REM output is regulated through episode frequency rather than duration.

## 1. Introduction

The alternation between non-rapid eye movement (NREM) and rapid eye movement (REM) sleep is one of the most prominent ultradian rhythms in mammalian physiology (Dement and Kleitman, 1957; McCarley and Hobson, 1975), yet the mechanism governing its timing and regulation remains unsettled after four decades of inquiry.

That a REM episode is followed by an interval whose length reflects the duration of that episode is an old observation, and it long predates the theories built on it. It was reported in monkeys (Weitzman et al., 1965), in cats (Ursin, 1970), and subsequently in rats (Zamboni et al., 1999), in humans (Barbato and Wehr, 1998; Cajochen et al., 2024), and in mice by Le Bon et al. (2007) — the cohort reanalysed here — and later independently by Weber et al. (2018). The regularity itself has never been seriously disputed. What it means has been.

The modern debate was crystallised in 1994, when two influential accounts read the same regularity in opposite directions, locating the primary regulatory variable on opposite sides of the REM episode. Benington and Heller (1994) proposed an NREM-driven framework in which REM propensity accumulates homeostatically during NREM sleep; on this view the intervening NREM interval is an accumulating drive that triggers the next REM episode once a threshold is reached, implying that longer NREM intervals should culminate in longer or more intense REM episodes. They observed the forward correlation in rats and interpreted it as evidence that REM sleep acts to reset or sustain subsequent NREM sleep, describing REM as a prerequisite that permits NREM to continue. Concurrently, Vivaldi et al. (1994) proposed a REM-instigated framework, in which the duration of a REM episode governs the duration of the subsequent inter-REM interval (IREM) while the interval does not feed forward to determine the next REM episode. The 1994 divide was therefore not about whether the association exists, but about which side of the REM episode carries the controlling variable.

Treating the preceding REM episode as the continuous determinant of the entire subsequent interval nevertheless faces a major empirical obstacle: the extreme variability and wake fragmentation of inter-REM intervals, particularly in polyphasic rodents and during the active dark phase. In 2020, two independent lines of reasoning converged on the idea of a post-REM refractory process. Le Bon (2020), from a theoretical reassessment of NREM-REM relationships, proposed that REM may impose a refractory constraint on its own recurrence, while Ocampo-Garcés et al. (2020), using an experimental REM-deprivation/permission paradigm in rats, demonstrated a transient REM-dependent reduction in REM transitions and explicitly proposed a short-term REM refractory period. The Asymmetrical Hypothesis subsequently formalized this view (Le Bon, 2021): preceding REM was proposed to impose a Post-REM Refractory Period (PRRP), followed by a Permissive Period (PP; originally Lambda) in which REM onset becomes increasingly dependent on other physiological and behavioral influences.

The critical empirical question is therefore not whether REMpre and IREM are correlated, but how long information carried by REMpre remains detectable, and what component of the interval discharges the constraint.

Subsequent propensity analyses have described early post-REM suppression and its temporal evolution increasingly directly (Park et al., 2021; Ginsberg et al., 2024; Akhavan et al., 2026). An early trough that shifts with REMpre can, however, be produced both by a sharply bounded refractory mechanism and by a steep but continuous homeostatic process. The present analysis therefore separates three quantities that should not be conflated: the lower envelope of earliest recurrence, the duration of strong suppression and recovery, and the later horizon at which REMpre ceases to provide detectable information.

We addressed four questions: whether the relation is directionally asymmetric; whether post-REM suppression is better described as continuous determination of the whole interval or as a finite-memory regime; whether accumulated NREM is a better predictive currency than elapsed time, particularly when wake fragments the interval; and how closely spaced episodes and 24-hour output are organized when episode duration is itself constrained.

Here, these questions were tested in continuous 24-hour polysomnographic recordings from 25 unperturbed mice across three genetic backgrounds (129Sv/Ev, CD1, and C57BL/6). Of the 1,968 consecutive inter-REM intervals in the 25 recordings, 1,955 met the validity criterion (Methods).

## 2. Materials and Methods

### 2.1 Animals and Polysomnographic Recordings

Continuous 24-hour polysomnographic recordings (EEG/EMG) were analyzed from 25 adult male mice belonging to three strains: 129Sv/Ev (n=10), CD1 (n=8), and C57BL/6 (n=7). The recordings form a subset of a previously described cohort (Le Bon et al., 2007); of the original 26 animals, one recording could not be identified unambiguously against the laboratory records and was excluded. Animals were housed individually under a 12:12-hour light/dark cycle (lights on at 07:00 h, ZT0) with food and water ad libitum at 24 ± 1°C. The animal protocol was conducted under French authorisation and complied with the Principles of Animal Care (NIH publication 86-23, revised 1985); permission for the present reuse was obtained from the host laboratory.

Vigilance states were scored offline in contiguous 15-s epochs as Wakefulness, Slow-Wave Sleep (SWS1 and SWS2, combined here as NREM), or REM sleep, according to the original study criteria; most recordings contained the expected 5,760 epochs. Because several archived files contained short missing segments or timestamp discontinuities, temporal gaps were identified explicitly and intervals crossing a gap or unscored segment were censored, reducing the dataset from 1,968 to 1,955. Analyses of 24-hour totals were additionally checked in near-complete recordings.

### 2.2 Interval and Episode Definitions

A REM episode was a run of consecutive 15-s epochs scored as REM, its duration being epochs × 0.25 min; the inter-REM interval (IREM) ran from the offset of one episode (REMpre) to the onset of the next (REMpost), and its elapsed duration, accumulated NREM and accumulated wake were computed separately. The 25 recordings contained 1,993 REM episodes. Phase was assigned as Light or Dark by clock time. Closely spaced episodes were analysed across a range of preceding separations rather than at a single post hoc sequential-cycle threshold.

### 2.3 Statistical Modeling and Guardrails Against Tautology

Distributional diagnostics guided the choice of estimators. All three interval variables are strongly right-skewed (skewness 1.16, 2.36 and 4.82 for preceding REM duration, accumulated NREM and elapsed interval duration; excess kurtosis 0.96, 9.41 and 38.80).

Pearson coefficients proved unstable under this skew: removing the upper 1% of accumulated NREM shifted the forward Pearson coefficient by +0.044 (0.416 to 0.460) against -0.003 for the rank coefficient (0.506 to 0.504), so bivariate associations are reported as Spearman correlations. Mixed-effects models were fitted on the log-log scale, where residuals are approximately symmetric (skewness -0.40, excess kurtosis 0.73) rather than markedly non-normal on the raw scale (2.70 and 12.46); the coefficient is then an elasticity, directly comparable with published log-log slopes. Raw-scale slopes are reported descriptively, with confidence intervals obtained by resampling mice rather than intervals. The quantile-regression and hazard analyses that carry the principal results make no distributional assumption.

Repeated cycles were treated as clustered within mouse. Directional associations were evaluated descriptively and with linear mixed-effects models; the principal REMpre→NREM model included random intercepts and random slopes by mouse. Mouse-by-mouse correlations were also reported. The lower envelope was estimated by quantile regression at the 5th percentile. Linear through-origin, linear with intercept, power-law and quadratic forms were compared by leave-one-mouse-out quantile loss, and binned percentiles were bootstrapped by resampling mice. To avoid in-sample definition of a putative boundary, k was re-estimated in leave-one-mouse-out folds and tested in the held-out mouse. Episode specificity was evaluated with 10,000 within-mouse permutations of REMpre pairing. Observations exactly on a boundary were not treated as violations. REM recurrence was analysed prospectively using discrete-time hazard models. REMpre effects were estimated as a function of accumulated NREM, with cluster bootstrap resampling at the mouse level (5,000 resamples). A segmented recovery model was compared with smooth alternatives to estimate whether a localized change in recovery slope was supported. The predictive horizon was defined operationally as the region after which the bootstrap interval for the REMpre effect included zero and no sustained REMpre effect re-emerged. To compare candidate currencies, otherwise equivalent prospective hazard models were indexed either by accumulated NREM or by elapsed chronological time and evaluated by grouped cross-validation with mice held out from model fitting. For closely spaced REM episodes, episode morphology was compared with isolated episodes and the contribution of the preceding REM episode was tested over a range of separation thresholds. REM episode termination was characterized by Weibull survival models. Macro-level analyses avoided regressions containing algebraically coupled quantities (REMtot=N×mean REM); instead, variability in episode number and mean episode duration was compared on the log scale, with sensitivity analyses restricted to near-complete recordings.

## 3. Results

### 3.1 Macro-Architecture and Sleep Parameters

Summary sleep parameters are shown in Table 1. Interval-level analyses use the gap-censored dataset; the 24-hour totals in Table 1 retain their descriptive role.

**Table 1.** Summary 24-hour sleep parameters across the 25 recordings (mean ± SD, median, and range).

| Parameter | Mean ± SD | Median | Range (Min – Max) |
| --- | --- | --- | --- |
| Total Sleep Time (TST, min) | 728.2 ± 86.8 | 724.2 | 528.2 – 903.8 |
| Total NREM Time (min) | 643.4 ± 80.3 | 637.2 | 475.2 – 831.0 |
| Total REM Time (min) | 84.8 ± 16.4 | 83.0 | 53.0 – 117.8 |
| Total Wake Time (min) | 706.5 ± 87.5 | 710.5 | 535.8 – 911.8 |
| Total Number of Cycles / Episodes (N) | 79.7 ± 22.0 | 74.0 | 49.0 – 127.0 |
| Mean REM Episode Duration (min) | 1.09 ± 0.15 | 1.09 | 0.88 – 1.43 |

**Table 2.** Quantile regression of accumulated NREM on preceding REM episode duration, across conditional quantiles. The through-origin coefficient k is the boundary estimate constrained to pass through zero.

| Quantile ($\tau$) | Intercept (min) | Slope ($\beta$, min/min) | Through-origin $k$ | Interpretation |
| --- | --- | --- | --- | --- |
| 0.05 | -0.300 | 1.400 | 1.000 | Primary lower-envelope estimate |
| 0.10 | -0.400 | 2.600 | 2.300 | Lower tail |
| 0.25 | 0.333 | 3.333 | 3.600 |  |
| 0.50 | 1.719 | 4.125 | 5.333 | Median |
| 0.75 | 4.875 | 4.500 | 8.500 |  |
| 0.90 | 9.750 | 5.000 | 14.500 | Upper tail |

Because intervals were re-extracted from the raw epoch files rather than reused from the original derived tables, the published values of Le Bon et al. (2007) provided an external check on the pipeline. Under the original episode definitions the animal-level correlations were recovered closely: cycle number with total REM 0.896 (published 0.907) and with total NREM 0.151, non-significant (published 0.236, non-significant). At the cycle level, the rank correlation between REM episode duration and the NREM content of the following interval was 0.520 (published 0.520), and with the duration of that interval 0.463 (0.451), while both backward associations remained null. Rank correlations are reported here because both distributions are strongly right-skewed. Total NREM tallies are higher in the present analysis because it covers the full 24 h, whereas the original tallies excluded incomplete and light-dark transition cycles; one of the original 26 animals was not available (Methods). The dataset therefore reproduces the published description of this cohort, the baseline against which the analyses below should be read.

### 3.2 Adjudicating the 1994 Divide: Directional Asymmetry

The directional asymmetry was reported for this cohort originally (Le Bon et al., 2007) and is recovered here as a starting point rather than a new finding. Accumulated NREM showed no meaningful relation with the duration of the subsequent REM episode (Spearman rho=0.015, p=0.50), consistent with the absence of NREM-to-REM duration scaling. By contrast, REMpre robustly predicted subsequent accumulated NREM (Spearman rho=0.506, p<10^-100). The association with elapsed interval duration was weaker (rho=0.453), and successive REM episode durations remained uncorrelated (rho=0.003, p=0.90).

A random-intercept/random-slope mixed model fitted on the log-log scale returned an elasticity of 0.562 ± 0.033 (95% CI 0.497–0.627; p<10^-60), substantially steeper than the log-log slopes of approximately 0.33 previously reported in humans and rodents. On the raw scale, and reported descriptively, each additional minute of REMpre was followed by approximately 3.9 min of additional NREM (95% CI by mouse-level resampling 3.51–4.35). The result was not produced by between-animal pooling: all 25 mice showed positive within-animal associations, with mean Spearman rho=0.462 ± 0.136 (range 0.202–0.667); 24 of 25 were individually significant. The forward association is therefore a reproducible within-animal property of the recordings rather than a consequence of combining animals with different sleep architectures — a decomposition that the original pooled analysis could not provide.

Genetic background did not materially alter the REMpre→NREM relationship; strain contrasts were non-significant (p=0.49 and p=0.66). The directional result was therefore retained as a common effect across the three strains.

### 3.3 A REMpre-Dependent Lower Envelope, Not an Absolute Floor

The lower tail of the conditional NREM distribution was used to test whether preceding REM imposes an early recurrence constraint. The 5th-percentile boundary is not linear in REMpre. Expressed as a through-origin coefficient it is close to k=1.0 min accumulated NREM per min REMpre, and leave-one-mouse-out folds centred tightly on that value (mean approximately 1.03); but it is a linear approximation to a convex boundary. In the same cross-validation a quadratic boundary outperformed the linear one (quantile loss 697.7 against 719.2, a 3.0% improvement) and outperformed power-law and linear-with-intercept alternatives; fitted through the origin, the quadratic term was highly significant while the linear term was not (p<0.001 and p=0.069). The curvature is not an artefact of sequential cycles, which are concentrated at short REMpre (24% of intervals shorter than 2 min follow episodes below 0.6 min, against none above 2.4 min): excluding intervals shorter than 2 or 3 min, or those with less than 1 min of accumulated NREM, leaves the quadratic coefficient positive and the quadratic form preferred in cross-validation, although its magnitude falls from 0.88 to between 0.33 and 0.47. The boundary therefore rises faster than the preceding episode rather than scaling proportionally with it: on the full dataset the fitted boundary rises from approximately 0.5 min of NREM after a 0.5-min episode to approximately 6 min after a 3-min episode. Because only 143 intervals follow episodes longer than 2.4 min, the curvature is established more securely than its exact functional form (Figure 1A).

**Figure 1.**
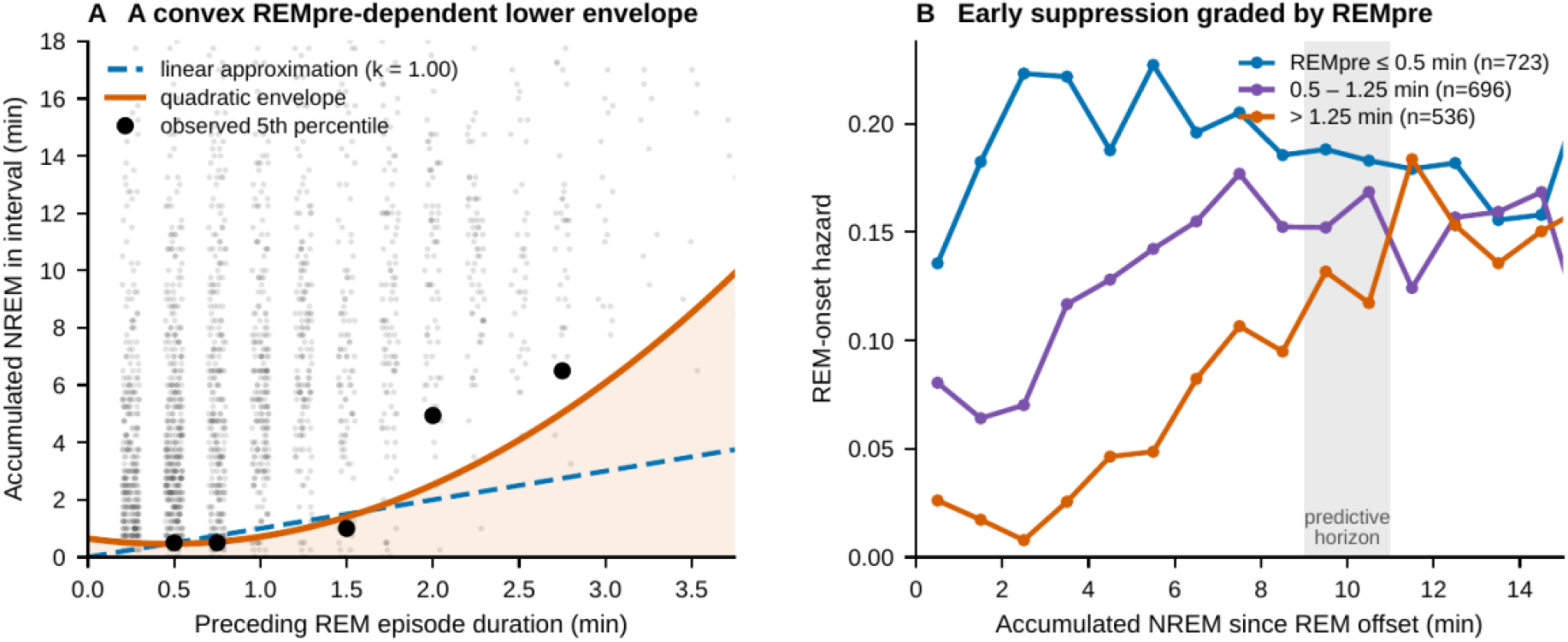
Post-REM interval architecture in 25 mice (1,955 valid inter-REM intervals). (A) A convex REMpre-dependent lower envelope: accumulated NREM to the next REM episode plotted against preceding REM episode duration. Black points are the observed fifth percentile within bins of REMpre; the orange curve is the cross-validated quadratic boundary and the dashed blue line its linear through-origin approximation (k = 1.00). The boundary rises faster than REMpre rather than scaling proportionally with it. (B) Empirical REM-onset hazard as a function of accumulated NREM, stratified by tertile of preceding REM duration: early suppression is graded by REMpre (0.14, 0.08 and 0.03 in the first minute) and the three curves converge across the shaded predictive horizon at 9-11 min.

Across held-out mice, 119 of 1,955 intervals (6.09%) fell below the fold-specific prediction. Under 10,000 within-mouse permutations the rate averaged 10.20 ± 0.39% (range 8.70–11.82%), and no permuted dataset reached a value as low as observed (empirical p≈0.0001 with the usual +1 correction). Early recurrence is therefore structured by the specific duration of the preceding episode. Because approximately 6% of observations nevertheless fall below the boundary, this is better read as a robust REMpre-dependent lower envelope than as an impenetrable physiological floor.

The finding did not depend on a single operational REM definition: allowing brief wake or non-REM interruptions shifted the coefficient upward rather than abolishing the boundary, and the most conservative strict-episode definition is retained here.

Higher conditional quantiles showed progressively larger slopes, indicating that dispersion increases above the early envelope. The envelope coefficient cannot therefore be read as the duration of the entire post-REM process; it separates the earliest recurrence boundary from the more gradual decay tested below.

### 3.4 Post-REM Recovery and a Finite Predictive Horizon

Prospective discrete-time hazard analysis showed marked REMpre-dependent suppression early after REM. Per additional minute of REMpre, the odds ratio for recurrence was approximately 0.27 in the first minute of accumulated NREM, 0.14 at 1–2 min and 0.10 at 2–3 min, then attenuated progressively (0.35 at 4–5, 0.52 at 6–7, 0.68 at 8–9 and 0.79 at 9–10 min). In 5,000 mouse-level bootstrap resamples, the REMpre coefficient remained significantly negative through 9–10 min (beta=-0.232; bootstrap 95% CI -0.468 to -0.005) but included zero at 10–11 min (beta=-0.226; -0.509 to +0.026) and thereafter. We therefore estimate a predictive horizon of approximately 10 min accumulated NREM (Figure 1B), beyond which preceding REM duration no longer provides a sustained detectable contribution to recurrence probability.

### 3.5 Accumulated NREM as the Better Predictive Currency

Wake fragmentation provided a natural test of chronological time against accumulated NREM. In Light (n=1,283; mean intervening wake 5.4 min), REMpre correlated with accumulated NREM at rho=0.514 and with elapsed IREM at rho=0.475; in Dark (n=672; mean intervening wake 13.2 min), the NREM association was preserved at rho=0.497 whereas the elapsed-time association fell to rho=0.426. The same distinction held prospectively. In five-fold grouped cross-validation, with five mice held out per fold, an accumulated-NREM hazard model achieved a cumulative predictive log-likelihood of -8472.17 compared with -8504.59 for an otherwise equivalent elapsed-time model (Δ log-likelihood +32.41 in favour of NREM). The NREM-indexed model won in four of five folds; the remaining fold was nearly tied. The aggregate difference clearly favours the NREM-indexed model, but it accrues across 1,955 held-out intervals and corresponds to roughly 0.017 nats per interval: the advantage is consistent rather than large per observation. A simpler leave-one-mouse-out regression comparison likewise favored accumulated NREM (out-of-sample R^2^≈0.173) over elapsed IREM (≈0.042). These results support accumulated NREM as the more informative currency of the post-REM process, without implying that NREM accumulation alone causally determines REM recurrence.

### 3.6 Sequential and Single Cycles: A Graded Boundary Rather Than Two Regimes

The recent propensity literature separates sequential cycles — brief, NREM-poor inter-REM intervals — from single cycles, and fits two-component mixtures to the interval distribution in order to do so. Whether this reflects two regulatory regimes or a convenient cut through a single continuous distribution has not been tested directly. Two observations argue against two regimes. First, the REM episodes making up sequential cycles are not fragments of one interrupted episode: their individual durations match those of isolated episodes (median 1.00 min in both), whereas their summed duration does not. They behave as ordinary episodes that happen to occur close together, although morphology alone cannot establish biological independence.

Second, the boundary is graded rather than categorical. Adding the earlier REM episode to a mixed model predicting subsequent accumulated NREM improved fit at short separations (ΔAIC -3.13 at ≤1 min, -4.44 at ≤1.5 min, -8.71 at ≤2 min, -3.52 at ≤3 min, -6.24 at ≤4 min), weakened at ≤5 min (-0.72) and disappeared by ≤6 min (+1.67); across the unselected dataset the earlier episode added no information (ΔAIC +2.00, p=0.99). The contribution of an earlier episode therefore fades over roughly four to five minutes rather than switching off at a fixed threshold. The sequential-versus-single distinction is thus better read as the visible trace of a finite memory in the post-REM constraint than as evidence of two distinct state generators. This does not make the classification useless as a description, but a fixed cut-off approximates a graded property.

### 3.7 REM Episode Duration Is Strongly Constrained

Across the 25 animals, mean REM episode duration varied substantially less than episode number. At the episode level, no REM episode exceeded 4.25 min, whereas valid inter-REM intervals extended to 376.75 min.

A Weibull fit to REM episode duration yielded a shape parameter of approximately 1.477, clearly above unity, whereas the corresponding inter-REM shape was approximately 0.836 (Figure 2A). Thus, the conditional probability of REM termination increases as an episode progresses, unlike the much more dispersed intervals separating episodes. This supports an intrinsic duration constraint on REM episodes, but not a mathematically hard upper ceiling: a Weibull distribution with shape >1 remains unbounded.

**Figure 2.**
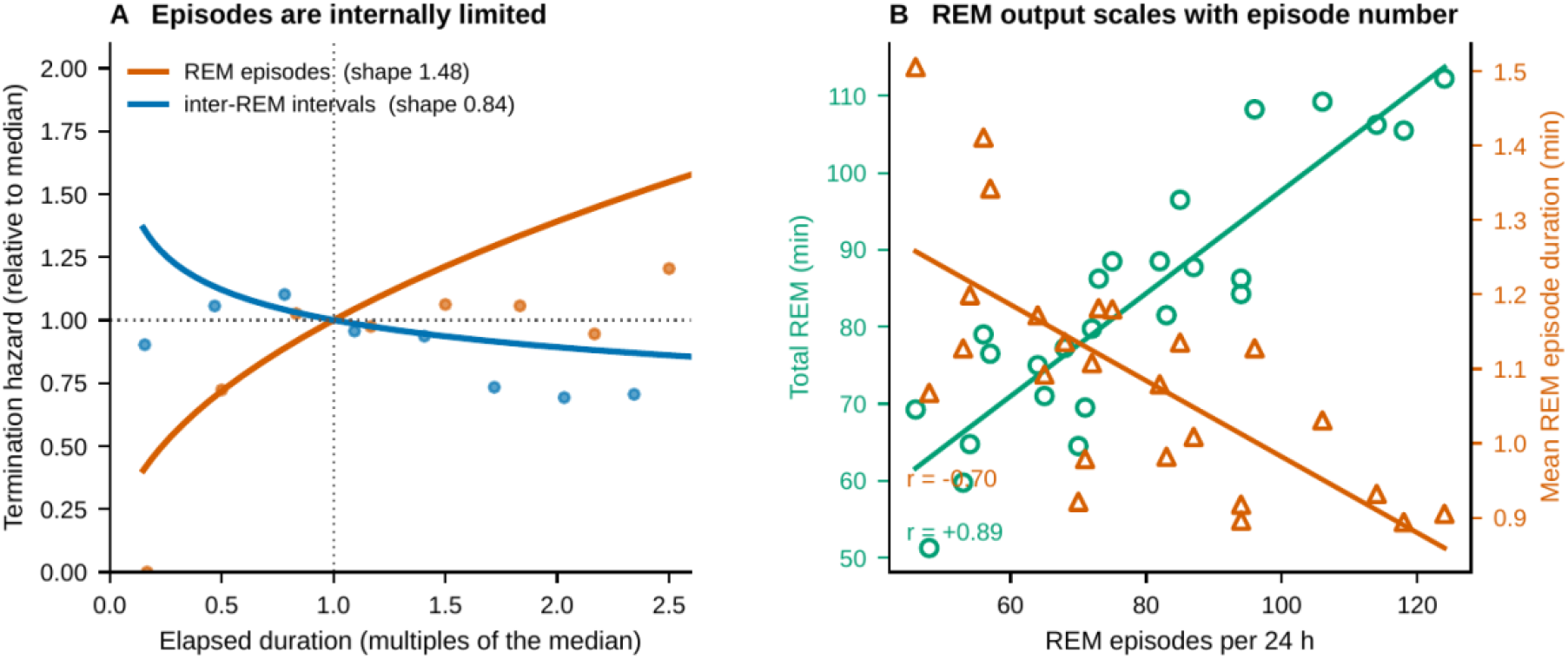
Episode-level and macro-level organization. (A) Termination hazard of REM episodes and of inter-REM intervals, each expressed relative to its own median duration; curves are fitted Weibull hazards, points are empirical estimates. The hazard of ending a REM episode rises with elapsed duration (shape 1.48) whereas that of ending an interval falls (shape 0.84), indicating that episodes are internally limited while intervals are not. (B) At the animal level, total 24-hour REM increases with episode number (green circles, r = +0.89) while mean episode duration decreases (orange triangles, r = -0.70); inter-animal variance on the log scale is about fourfold greater for episode number than for mean duration.

The term ‘quantum occurrence’ is therefore used, if at all, descriptively: REM is expressed in discrete episodes whose duration is relatively constrained, while the number of occurrences is considerably more variable. It should not be read as implying a literal fixed quantum or an analogy with physics.

### 3.8 Macro-Organization of REM Output

At the subject level, total 24-hour REM remained strongly associated with episode number (r≈0.90), whereas its association with mean episode duration was weak and non-significant (r=-0.34, p=0.10). Mean episode duration was, by contrast, clearly and inversely associated with episode number itself (r=-0.70, p<0.001; Figure 2B): animals with more episodes had shorter ones, yet total REM did not track episode length. The first was already reported for this cohort (Le Bon et al., 2007, r=0.907) and is not offered as new. These correlations are partly coupled by the identity REMtot=N×mean REM and cannot by themselves establish a regulatory mechanism.

Macro-organization was therefore evaluated by decomposition on the log scale, a regression of mean duration on REMtot being tautological. Variance of log episode number was 0.0741 compared with 0.0184 for log mean REM duration, a ratio of 4.0. In the 22 near-complete recordings, total REM remained strongly associated with episode number (r=0.868), whereas the association with mean duration was small (r=-0.152, p=0.50). Inter-animal variation in REM output is therefore expressed predominantly through variation in episode number rather than mean episode duration (Figure 2B).

## 4. Discussion

The reanalysis supports a robust asymmetry in ultradian REM organization but modifies the strongest version of the original Asymmetrical Hypothesis. A low-quantile recurrence boundary should not be equated with the full duration of a strictly bounded PRRP. Three empirically distinct features emerge instead: a convex REMpre-dependent lower envelope, a several-minute phase of strong suppression and recovery, and a finite predictive horizon near 10 min accumulated NREM. This reconciles the historical REMpre-IREM correlation with the extreme variability of the later interval.

### 4.1 The 1994 Divide Revisited: Direction Is Asymmetric

On an accumulation account, the NREM that precedes a REM episode should set how much REM follows. It does not: across 1,955 intervals, accumulated NREM was unrelated to the duration of the next REM episode. This is not a refutation of Benington and Heller (1994), who reported the same independence themselves. It does, however, remove episode size from the quantities that accumulated NREM can be said to govern, and leaves their account resting on timing alone.

By contrast, the forward REMpre→NREM relation was strong, reproducible, and present within every animal. The random-slope analysis is important here because it shows that the historical correlation is not generated by pooling mice with different characteristic REM durations or sleep amounts. In this limited directional sense, the data strongly favor an organization in which information is carried forward from a REM episode into the interval that follows.

Direction and currency should nevertheless be kept conceptually separate. REMpre carries information about the subsequent constraint, whereas the component of the interval that best tracks that constraint is accumulated NREM. The present findings therefore combine the forward direction emphasized by Vivaldi et al. with the importance of NREM emphasized by Benington and Heller, without requiring that NREM determine the size of the next REM episode.

### 4.2 Lower Envelope, Refractory Regime, and Predictive Horizon

The principal conceptual revision concerns the meaning of the PRRP. The lower envelope should not be equated with the duration of the refractory process. It identifies the earliest conditional boundary of REM recurrence, which at the 5th percentile is approximated by one minute of accumulated NREM per minute of REMpre but is in fact convex: the boundary rises faster than the episode, so that doubling the preceding episode more than doubles the NREM observed to intervene. Because held-out violations remain at approximately 6%, the data do not demonstrate an absolute h(t)=0 interval. They do, however, show that this lower tail is far more structured by the actual preceding REM episode than expected after within-mouse permutation. Hazard analysis reveals a broader refractory regime. REMpre exerts very strong suppression during the first several minutes of accumulated NREM, followed by progressive recovery. A segmented analysis in long-REM cycles placed the best bend in recovery near 6 min NREM, but the mouse-level bootstrap interval was broad (approximately 3–8 min) and the segmented model only modestly improved a single-slope alternative. The release is therefore better viewed as gradual than as a sharply localized switch. A third quantity can be estimated operationally rather than identified biologically: REMpre information persists after REM has become possible, but attenuates until it is no longer detectable under this model at approximately 10 min accumulated NREM. Thus, floor, PRRP duration, and predictive horizon are different estimands. The post-REM process is best described as a graded REMpre-dependent refractory regime with a strongly structured early boundary and finite memory, rather than as either a perfectly bounded window or continuous determination of the whole IREM.

The convexity is itself informative, and it revises the original formulation more than the linear coefficient does. A strictly proportional refractory period, as proposed in 2021, predicts a boundary linear in the preceding episode. The boundary is instead convex, and remains so when sequential cycles are excluded. Two readings are available: the process may saturate in its capacity to be discharged, or episodes reaching the upper part of the duration distribution may differ in kind from brief ones, in which case the boundary reflects a change in the episodes rather than in what follows them. The present data cannot separate these, but they exclude simple proportionality and make k a summary statistic rather than a parameter of the underlying process.

The shape of the pooled propensity function deserves comment, because it bears on how the curves reported since Park et al. (2021) should be read. Our pooled hazard reproduces the familiar rise-then-fall: it increases from 0.13 to 0.24 by approximately 11 min of accumulated NREM and declines thereafter. Part of that descending limb, however, requires no mechanism. Because REMpre sets a graded early boundary, intervals still at risk late are progressively enriched in long preceding REM episodes, which are precisely the intervals with the lowest early hazard: mean REMpre among survivors rose from 1.06 min at REM offset to 1.63 min at 15 min, and the proportion exceeding 1.25 min doubled from 27% to 54%. A mixture of exponentials matched to this stratum composition, with a constant hazard within each stratum and therefore no decline anywhere, nevertheless produced a pooled hazard falling from 0.19 at its maximum to 0.15 thereafter. A population-level decline can thus arise while no individual hazard declines, which is the standard consequence of unmodelled heterogeneity in survival data.

This does not establish that the observed decline is entirely artefactual. Normalizing time by each animal’s median interval, and additionally stratifying by REMpre tertile, attenuates but does not abolish it, so residual heterogeneity and a genuine late decline may both contribute. The implication is narrower and methodological: a non-monotonic pooled propensity function is not by itself evidence for circadian, arousal-related or other state-dependent modulation, and comparisons are better made on hazards conditioned on preceding REM duration and on animal identity. The same argument tempers our own predictive horizon: after approximately 10 min, REMpre may cease to separate the curves partly because selection has homogenized the surviving intervals with respect to REMpre, rather than because REM-history information has been fully discharged.

### 4.3 Accumulated NREM as the Currency of Post-REM Recovery

The Light-Dark dissociation and the held-out hazard comparison both favour accumulated NREM over elapsed time. Wake infiltration in Dark weakens the REMpre-elapsed-time relationship while leaving the REMpre-NREM relationship comparatively preserved, and, more importantly, accumulated NREM predicts recurrence better in mice excluded from model fitting. This is stronger evidence than an in-sample comparison, although it remains predictive rather than causal.

This is compatible with the observations of Weber et al. (2018), in which REM-off vlPAG GABAergic activity changes across post-REM sleep and wake. The correspondence is suggestive rather than a mechanistic identification, since the present recordings do not measure the circuit. We use ‘currency’ to mean the interval component that best indexes the decay of REM-history information, not proof that NREM is the sole causal discharge mechanism.

### 4.4 Closely Spaced Episodes Reveal Short-Lived Additivity, Not a Separate Regime

The sequential/single distinction is better treated as a continuum than as a taxonomy. Closely spaced bouts have episode durations comparable with isolated ones, and an earlier episode adds information about subsequent NREM accumulation only when the two occur within a few minutes, sensitivity across thresholds placing this residual additivity at approximately 4–5 min and none by about 6 min or in the unselected dataset. This short memory is compatible with residual post-REM inhibition carrying across a nearby episode, but does not prove a summation circuit.

### 4.5 REM Output Is Organized Predominantly Through Episode Occurrence

The macro-level findings complement the interval-level architecture. REM episode termination hazard increases as an episode progresses, while mean episode duration varies much less across animals than episode number. After removing the algebraically coupled regression, variance decomposition still shows approximately fourfold greater variability in episode number on the log scale. Thus, differences in 24-hour REM output are expressed predominantly through how often REM episodes occur rather than through large changes in their mean duration. Episode duration is constrained twice over: from within, by a termination hazard that rises as the episode progresses, and from without, by a recurrence boundary that rises faster than the episode itself. The second of these comes from the present data. Because the envelope is convex, the NREM accumulated before the next REM episode grows faster than the episode that preceded it, so that a given amount of REM distributed over few long episodes is accompanied by more intervening NREM than the same amount distributed over many short ones. This is an accounting relation rather than a mechanism: the data show what intervenes, not that it is required, and they say nothing about what ends an episode. The relation is nevertheless the one an architecture built on episode number rather than episode length would display. For the internal constraint — what actually ends an episode — one functional rationale is available, untested here. On Schmidt’s (2014) energy-allocation account, REM entails suspension of thermoregulatory defence, so a rising termination hazard is expected rather than incidental. The account predicts that the shape parameter should vary with ambient temperature, which recordings made at a single temperature cannot address. ‘Quantum occurrence’ is used here only as a descriptive shorthand for this organization: REM is expressed as discrete, duration-constrained events whose frequency is substantially more variable than their mean size; it does not imply a literal fixed quantum or an exclusive homeostatic mechanism. This macro-level organization links cautiously to the post-REM results. Each REM episode is not only duration-constrained; it also resets the local temporal context by initiating a new REMpre-dependent regime, so that increasing output through additional episodes creates additional post-REM constraints. Two complementary levels therefore emerge: at the ultradian level each episode transiently constrains the probability of the next occurrence, and at the 24-hour level variation in output is expressed through the number of these constrained events. The present observational data cannot establish the mechanism that selects episode number.

This two-level structure maps onto a distinction already drawn in the literature. Franken (2002) separated the short-term process governing REM timing from the long-term process governing REM amount over 24 h and beyond, and Ocampo-Garcés et al. (2020) likewise distinguished a long-term hourglass controlling REM recovery from a short-term process shaping ultradian organization. The present data specify what the long-term level operates on: variation in 24-hour REM amount is expressed through the number of episodes rather than through their duration, while the short-term process constrains when the next episode can occur. On this reading the two are not competing accounts of the same quantity but act on different ones — the interval and the count — which may explain how REM amount can be regulated across days while the timing of any individual interval remains highly variable.

### 4.6 Implications for the Concept of the Ultradian Sleep Cycle

This two-level organization also changes how the ultradian sleep cycle can be conceptualized. Consolidated human nocturnal sleep visually suggests a relatively regular sequence, whereas polyphasic mammalian sleep contains highly variable wake and NREM intervals between REM bouts; the familiar ‘NREM-REM cycle’ need not therefore be a unitary oscillator. The present findings weaken the need to treat the entire inter-REM interval as a single continuously determined cycle. A local REM-history effect is clearly present, but it is strongest early, relaxes over several minutes, and becomes non-detectable after a finite horizon. The robust REMpre-IREM association can thus arise from a local post-REM constraint embedded within a much more variable interval, so that later recurrence varies widely without contradicting the forward association. This is particularly relevant in polyphasic rodents, where wake and isolated NREM bouts make a visually regular cycle difficult to identify.

The data do not prove that no dedicated ultradian oscillator exists. They show instead that the observed interval architecture does not require preceding REM to determine the entire time to the next REM episode. An episodic asymmetrical account—REM followed by a finite-memory refractory regime embedded in a variable physiological and behavioral background—provides a parsimonious structural description of these recordings. Whether the same decomposition explains compact human nocturnal sleep remains an empirical question.

### 4.7 Limitations and Methodological Considerations

Several limitations constrain interpretation. These are observational recordings from an archived cohort; they cannot establish that transition hazard is literally zero at any early time, nor identify the causal neural substrate of recovery. The recovery bend near 6 min is imprecisely localized and is better regarded as evidence for gradual release than for a sharp change-point, and the predictive horizon, estimated from mouse-level bootstrap inference, should be replicated prospectively. A minority of recordings contained missing segments; intervals crossing gaps were censored and macro-level conclusions checked in near-complete recordings, but future datasets should be prospectively quality-controlled. Macro-level identities among total REM, episode number and mean duration impose unavoidable mathematical coupling, so the analysis emphasizes variance structure rather than causal interpretation. Finally, extension to other species, particularly humans with much longer REM episodes and more consolidated sleep, is necessary before claiming a universal architecture.

## 5. Conclusions

Across 25 mice, the organization of REM recurrence is strongly asymmetrical. Preceding REM duration predicts subsequent NREM accumulation within every animal, whereas accumulated NREM does not scale the duration of the next REM episode. The post-REM effect is not adequately described either as continuous determination of the whole interval or as a refractory period whose duration is identical to a strict lower floor. Instead, the data identify a convex REMpre-dependent lower envelope, strong suppression that recovers over several minutes of accumulated NREM, and a finite predictive horizon of approximately 10 min. Accumulated NREM predicts this process better than chronological time, particularly when wake fragments the interval. Closely spaced REM episodes retain short-lived additive information. REM episodes are themselves duration-constrained, and inter-animal variation in 24-hour REM output is expressed predominantly through episode number rather than episode length. Two linked levels of organization therefore emerge: at the ultradian level each episode initiates a transient, REM-duration-dependent refractory regime, and at the 24-hour level output varies through how many such episodes occur. The revised Asymmetrical Hypothesis is best framed as regulation of discrete REM occurrences embedded in a graded, NREM-indexed refractory architecture with an early boundary, finite REM-history memory, and a later multifactorial permissive period.

## Data Availability Statement

The epoch-level scoring of all 25 recordings, the derived interval dataset, and the analysis code reproducing every reported value and both figures are deposited on Zenodo (doi: 10.5281/zenodo.22688770). Reuse of the original recordings was authorised by the host laboratory.

## Ethics Statement

No new animal experimentation was performed for this study. The recordings analysed here are a subset of a previously published cohort (Le Bon et al., 2007), obtained under French regulations then in force and in compliance with the Principles of Animal Care (NIH publication 86-23, revised 1985). Permission for the present reanalysis was obtained from the host laboratory. The original protocol identifier is no longer retrievable.

## Author Contributions

OLB: Conceptualization, Data curation, Investigation, Methodology, Project administration, Software, Validation, Writing – original draft, Writing – review and editing

## Generative AI Statement

The author used Anthropic’s Claude, OpenAI’s ChatGPT and Google’s Gemini as analytical and editorial assistants throughout this work. Claude wrote and executed the Python code used to reconstruct inter-REM intervals from the raw epoch files and to perform the analyses reported here — mixed-effects models, quantile regression, permutation and cross-validation procedures, discrete-time hazard models, Weibull survival fits and bootstrap resampling — produced the figures and contributed to drafting and revising the text. ChatGPT and Gemini were used to double-check at every step. The author conceived the study and its questions, supplied and verified the source recordings, specified or approved each analysis, checked the outputs against the published description of the same cohort, and identified and corrected errors in the assistant’s analyses and text. The author takes full responsibility for the content, the accuracy of the reported values and the conclusions drawn. The assistants are not authors and made no independent scientific judgement. The analysis code is deposited with the data (doi:10.5281/zenodo.22688770) so that every reported value can be recomputed independently.

